# Monitoring of group mates in relation to their activity in mandrills

**DOI:** 10.1101/810440

**Authors:** Gabriele Schino, Martina Scerbo

## Abstract

Primates are known to have considerable knowledge about the social relationships that link their group mates, and are likely to derive this information from observing the social interactions that occur in their social group. They may therefore be hypothesized to pay particular attention to the social interactions involving group mates. In this study, we evaluated how the attention captive mandrills (*Mandrillus sphinx*) devote to their group mates was modulated by the behavior of the latter. Mandrills looked most frequently at foraging individuals and least frequently at sleeping invividuals. Mandrills also looked at grooming individuals more than at individuals that were simply sitting in contact. Grooming dyads were looked at regardless of the social rank and kinship of the individuals involved. These results contribute to our understanding of how primates obtain their social knowledge.

## INTRODUCTION

In order to survive, primates need in-depth and updated knowledge about their environment. Such knowledge can be aquired either individually or socially, by observing how group companions interact with potential sources of food or danger (Cook & Mineka, 1990; Rapaport & Brown, 2008). In order to compete successfully, primates also need updated knowledge about the social relationships that link their group mates. Indeed, research conducted in the last 40 years has shown that primates have considerable social knowledge. Not only are they aware of relatively stable characteristics of third-party social relationships such as dominance and friendships (reviews in Seyfarth & Cheney, 2012, 2015), but they also appear to keep updated information on more transient phenomena such as sexual consortships, take-overs of one-male units, and postconflict reconciliation (Crockford et al., 2007; Judge & Bachmann, 2013; le Roux & Bergman, 2012).

Given that the verbal exchange of social information is obviously precluded to nonhuman primates, they must necessarily obtain their social information individually and observationally. We know very little, however, about what aspects of the social life of their group mates they attend to. In one of the few studies that addressed this issue, Tiddi et al. (2017) showed that Japanese macaques (*Macaca fuscata*) did not obtain information about the kinship relationships of their group mates by remembering early mother-infant interactions, but seemed to keep an updated record of affiliative interactions occurring among group mates. They apparently relied on a rule of thumb that considered as related all dyads that exchanged frequent affiliation, and were thus unable to distinguish the kin from the "friends" of their group mates.

If primates keep some sort of record of the interactions that occur among their group companions, we may hypothesize that they must pay particular attention to such interactions. The available information about how the attention primates pay to group mates is modulated by their behavior is scanty. A few studies showed primates (and non-primates) are interested in observing others engaged in ecologically relevant activities such as food manipulation or extraction (Ottoni et al., 2005; Range & Huber, 2007; Scheid et al., 2007). Barbary macaques (*M. sylvanus*) have been shown to attend to scratching in others, possibly as a way to monitor their emotional state (Whitehouse et al., 2016). Bonobos (*Pan paniscus*), showed an attentional bias towards images of sexual or grooming behavior (Kret et al., 2016). Overall, it is clear that this is a subject that requires further investigation.

In this study, we first replicated previous analyses of the effects of dominance rank and kinship on attention paid to group mates (Schino & Sciarretta, 2016). Then we evaluated how the attention paid by captive mandrills to their group mates is modulated by their activity. Given the obvious ecological relevance of finding food, we expected attention directed to foraging individuals to be high (although finding food may be less relevant in captivity). We hypothesized that mandrills would pay particular attention to the social interactions of group mates as a consequence of the need to monitor the state of their social relationships. We thus compared the attention paid to grooming individuals to that paid to individuals that are simply sitting in contact, as well as the attention paid to individuals that are in proximity to a group mate to that paid to lone individuals (see the Methods for details and justification). We also expected that active individuals would attract more attention. We thus compared the attention paid to sleeping and awake group mates. Exploratory analyses tested whether the effects of dominance rank and kinship was modulated by the activity of the target, and whether the attention paid to grooming was modulated by individual characteristics. Here, we hypothesized that rare grooming events should attract more attention than common grooming events.

## METHODS

### Ethical Note

This was a purely observational study conducted in a zoo setting. It complied with the Italian law, which requires no authorization for such studies, and with the American Society of Primatologists’ principles for the ethical treatment of non-human primates.

### Subjects and Housing

The subjects of this study belonged to a captive group of mandrills living in the Rome zoo (Bioparco). Initially, the group included two males and 10 females, but one male and one female died shortly after the beginning of data collection, and the few data that had been collected on them were discarded. Details on the group composition can be found in Table 1.

**Table 1.** Subjects of the study.

| **Name** | **Sex** | **Rank** | **Matriline** | **Observation sessions as subject** | **Observation sessions as target** |
| --- | --- | --- | --- | --- | --- |
| Bart | m | 1 | b | 302 | 374 |
| Beta | f | 10 | b | 442 | 262 |
| Blanca | f | 2 | b | 407 | 564 |
| Bunni | f | 3 | b | 463 | 403 |
| Genni | f | 7 | g | 508 | 552 |
| Giorgia | f | 8 | g | 499 | 485 |
| Giulia | f | 5 | g | 410 | 468 |
| Greta | f | 6 | g | 445 | 321 |
| Irene | f | 4 | g | 481 | 593 |
| Malinka | f | 9 | m | 401 | 336 |

The group lived in a 240 m^2^ outdoor enclosure connected with indoor rooms. The enclosure was enriched with ropes, trunks and perches. Mandrills were fed twice a day with vegetables, fruits, seeds and monkey chow. Seeds were often dispersed in the substrate, and mandrills spent a considerable amount of time searching for food. Water was available ad libitum.

Information on maternal kinship was derived by demographic records. Animals were arranged in a linear dominance hierarchy on the basis of unidirectional dyadic aggressions using the I&SI method as implemented in DomiCalc (de Silva et al., 2017; Schmid & de Vries, 2013). Further information on the subjects of this study can be found in Table 1.

### Data Collection

Data on glances were collected during “focal dyad” observations lasting three minutes. Each observation session had a subject and a target. The observer recorded all glances directed by the subject to the target. We collected data on a single target for two reasons: first, we felt it would increase the reliability of data collection (focusing on a single target is easier than having to take into account all possible targets); second, our stringent criteria about the visibility and activity of the target (see below) were impossible to be monitored continuously on all possible targets.

Glances were defined as “orienting the head towards the target”. Note that since data were not collected when the subject was engaged in any social interaction (see below), this definition excludes the sustained looking associated with grooming or threatening. Since in primates staring directly at a target is often interpreted as a threat, glances directed at group mates are generally very brief. This is the reason most previous studies on social attention measured rates rather than durations. We followed this tradition. Furthermore, being glances very brief, measuring their duration would be nearly impossible under the observational conditions of this study. Actual gaze direction (as revealed by eye movement) helped identify glances but was not always recognizable by the observer. Recording actual gaze direction in freely interacting animals is extremely difficult. That is why most previous studies resorted to definitions that included head orientation (see the Supporting Information in Allan & Hill 2017). It is reasonable to assume that measuring head orientation as an estimate of gaze direction introduces noise but not bias in the data.

Observation sessions were not initiated if an aggression (involving any member of the social group) had occurred in the previous 10 minutes. At the beginning of the observation session, the subject had to be awake and not engaged in any social interaction. The target had to be visible by the subject, involved in one of the activities described in Table 2, and had to be at least 1 m away from the subject.

**Table 2.** Behaviors of target considered in this study.

| **Behavior** | **Definition** | **N° of observation sessions** |
| --- | --- | --- |
| Awake alone | Sitting or standing with eyes open; no other individual is within 2 m | 508 |
| Foraging | Eating or searching in the substrate | 683 |
| Grooming | Careful picking and/or slow brushing of another monkey's fur using the hands and/or the mouth; if interrupted for more than 15 s it is considered terminated; both grooming given and grooming received are included | 1525 |
| Proximity | Being within 1 m of another individual (but not in contact) | 1101 |
| Sleeping alone | Sitting or lying with eyes closed; no other individual is within 2 m | 416 |
| Sitting in contact | Sitting in physical contact with another individual who is also sitting | 125 |

Observation sessions were interrupted if: the subject or the target were not any more visible to the observer; the target was not any more visible to the subject; the subject was involved in any social interaction; an aggression occurred in the group; the target changed its activity, including a change in the direction of grooming; the distance between the subject and the target decreased to less than 1 m. Observation sessions lasting less than 30 s were discarded. We were interested in (and analyzed) the effects of characteristics of the grooming dyad such as the difference in dominance rank between groomer and groomee. That is why we interrupted the observation session if the direction of grooming changed. Note also that: i) observation sessions were rather short (3 minutes) so that this sort of interruption occurred rarely; ii) there is no reason to suppose that interrupting observation sessions introduced any bias in the data.

A total of 4358 observation sessions were available for analysis (details in Tables 1 and 2). We were especially interested in the monitoring of the social interactions of group mates, and biased data collection accordingly (Table 2). Our ability to obtain data was however constrained by the relative frequencies of the different behaviors so that, for example, sitting in contact was underrepresented in our sample.

We also conducted group scans every 15 minutes (for a total of 2196 scans) recording all dyads engaged in grooming. Finally, we recorded aggressive events (threats, chases and physical assaults) ad libitum.

### Data analysis

All our analyses were within-subject (fixed effect) conditional Poisson regressions with bootstrap standard errors. Within-subject regressions allow the use of multiple data points per subject while avoiding pseudoreplication. In within-subject regressions, each individual is compared only with itself (similarly to a paired sample t test), so as to exclude the effects of unknown and unmeasured variables and avoid the confusion between within-subject and between-subject effects (Allison, 2009; van de Pol and Wright, 2009).

The dependent variable of all analyses was the count of glances recorded in each observation session (i.e., each observation session contributed one data point). The duration of the observation sessions was entered as an exposure variable. Independent variables included the rank of the target, the degree of maternal kinship between the subject and the target, and the activity of the target. The latter was entered as a dummy variable (i.e., as a set of six indicator variables each corresponding to one of the activities) and we assessed its significance using a Wald test that tested the null hypothesis that the effects of all the indicator variables were jointly zero.

Pairwise comparisons of the effects of the different activities of the target applied the Šidàk correction in order to control for repeated testing. Beside presenting all possible pairwise comparisons, we focused on specific pairs of activities comparing target behaviors that differed, as much as possible, in a single key aspect. In particular, we felt it important to compare behaviors that were comparable in terms of general activity level, as more active animals may simply be more conspicuous (see also below for a comparison focusing directly on activity level). “Foraging” was compared both with “Awake alone” and with “Proximity” (since our definition of Foraging did not specify whether the target should be alone or not). We compared “Grooming” with “Sitting in contact”, since both these contexts implied that two animals were stationary and very close to each other. We compared “Awake alone” with “Proximity” since for both contexts our definition did not specify whether the animals were sitting or standing and the only difference was the presence (within 1 m) of a group mate. Finally, we compared “Sleeping” with “Awake alone” since in both contexts no group mate was present nearby.

A separate analysis (again, a within-subject conditional Poisson regressions with bootstrap standard errors) focused on grooming (i.e., was limited to those observation sessions in which the target was involved in grooming). It tested the effects of characteristics of the grooming dyad such as their kinship, rank distance, and baseline frequency of grooming on glance rate.

All statistical analyses were run on Stata 14.2 (StataCorp, 2015).

## RESULTS

Both target rank and subject-target kinship affected the rate of glances directed by the subject to the target: mandrills directed more glances at high-ranking and at unrelated individuals (coeff.=−0.036, z=−8.22, N=4358, P<0.001 and coeff.=−0.262, z=−3.05, N=4358, P=0.002, respectively). Controlling for rank and kinship, the behavior of target affected potently the glance rate of the subject (Wald test: *χ*^2^=363.57, df=5, P<0.001; Fig. 1).

**Figure 1.**
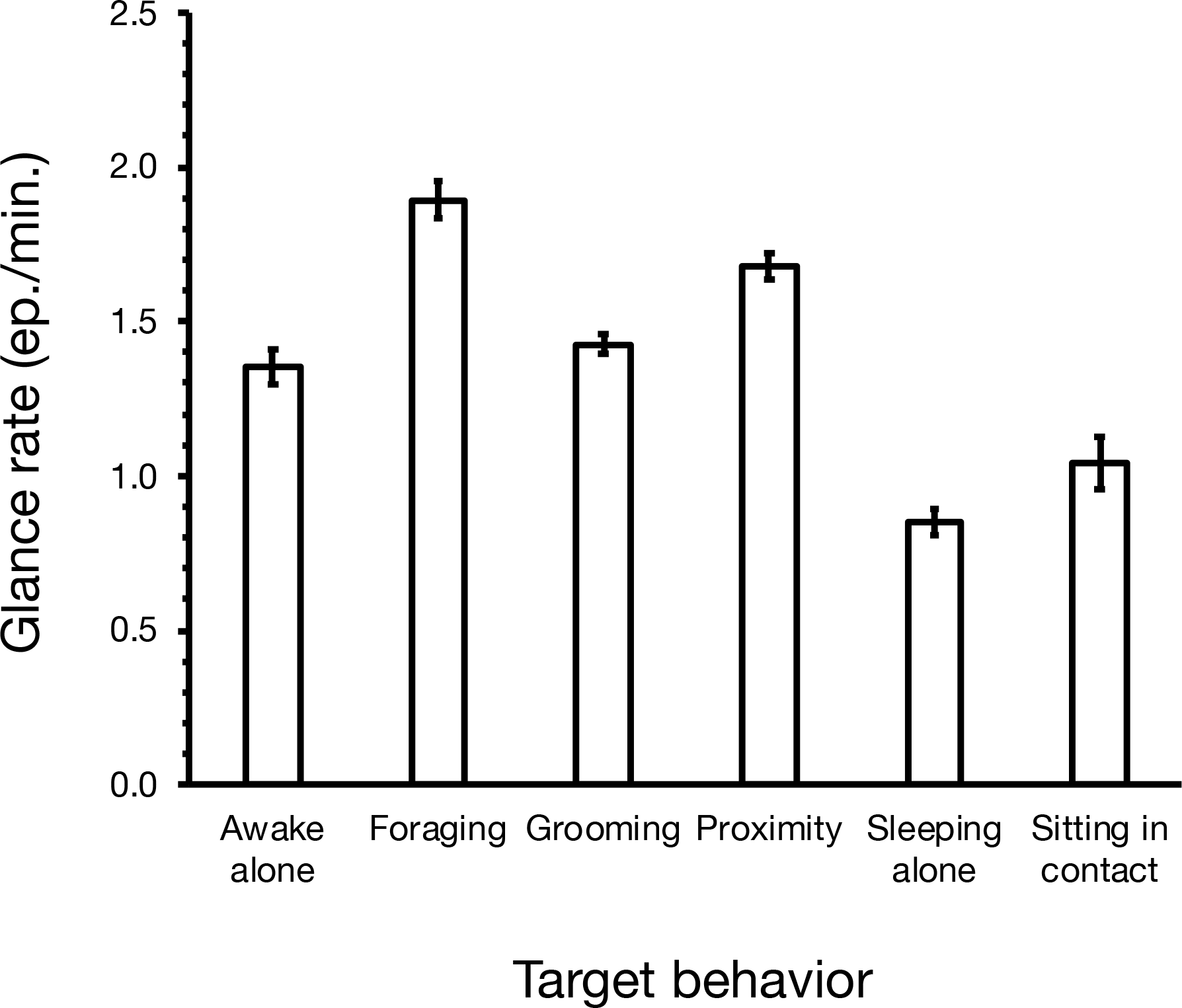
Effects of the target behavior on the rate of glances the subject directed to the target. Marginal means and standard errors, controlling for variation in the target dominance rank and in target-subject kinship.

Pairwise comparisons between target behaviors showed several significant differences (Table 3). Foraging was the target behavior most looked at by mandrills; specifically, it was looked at more than both “Awake alone” and “Proximity”. Confirming our predictions, animals engaged in grooming were looked at more than animals that were simply sitting in contact to each other. Similarly, animals that were in proximity to a group mate were looked at more frequently than lone animals (compare “Proximity” with “Awake alone”). Finally, awake individuals were looked at more than sleeping individuals (compare “Awake alone” with “Sleeping alone”).

**Table 3.** Glances directed at the target in relation to its behavior: pairwise comparisons.

|  | **Awake alone** | **Foraging** | **Grooming** | **Proximity** | **Sleeping alone** | **Sitting in contact** |
| --- | --- | --- | --- | --- | --- | --- |
| **Awake alone** | | $z=-4.77$<br>$P<0.001$ | $z=-1.01$<br>$P=0.997$ | $z=-3.94$<br>$P=0.001$ | $z=7.16$<br>$P<0.001$ | $z=4.53$<br>$P<0.001$ |
| **Foraging** | $z=4.77$<br>$P<0.001$ | | $z=7.23$<br>$P<0.001$ | $z=2.46$<br>$P=0.191$ | $z=14.87$<br>$P<0.001$ | $z=6.72$<br>$P<0.001$ |
| **Grooming** | $z=1.01$<br>$P=0.997$ | $z=-7.23$<br>$P<0.001$ | | $z=-8.14$<br>$P<0.001$ | $z=9.34$<br>$P<0.001$ | $z=5.10$<br>$p<0.001$ |
| **Proximity** | $z=3.94$<br>$P=0.001$ | $z=-2.46$<br>$P=0.191$ | $z=8.14$<br>$P<0.001$ | | $z=11.05$<br>$P<0.001$ | $z=7.71$<br>$P<0.001$ |
| **Sleeping alone** | $z=-7.16$<br>$P<0.001$ | $z=-14.87$<br>$P<0.001$ | $z=-9.34$<br>$P<0.001$ | $z=-11.05$<br>$P<0.001$ | | $z=-2.63$<br>$P=0.120$ |
| **Sitting in contact** | $z=-4.53$<br>$P<0.001$ | $z=-6.72$<br>$P<0.001$ | $z=-5.10$<br>$p<0.001$ | $z=-7.71$<br>$P<0.001$ | $z=2.63$<br>$P=0.120$ | |

In order to evaluate whether the effects of target rank and of subject-target kinship were modulated by the behavior of the target, we repeated the first analysis presented above adding the interactions between target rank and target behavior and between subject-target kinship and target behavior. The interaction between target rank and target behavior was significant (Wald test: *χ*^2^=60.38, df=5, P<0.001), while that between subject-target kinship and target behavior was not ((Wald test: *χ*^2^=6.85, df=5, P=0.232). Analyses of the effect of target rank split by target behavior revealed that glance rate increased with increasing target rank for all behaviors examined but for grooming and sitting in contact (Fig. 2). Note that while the nonsignificant effect obtained for targets engaged in sitting in contact may be explained by the relatively small sample size, this is not a likely explanation for grooming, that had by far the largest sample size of all the activities we considered (Table 1).

**Figure 2.**
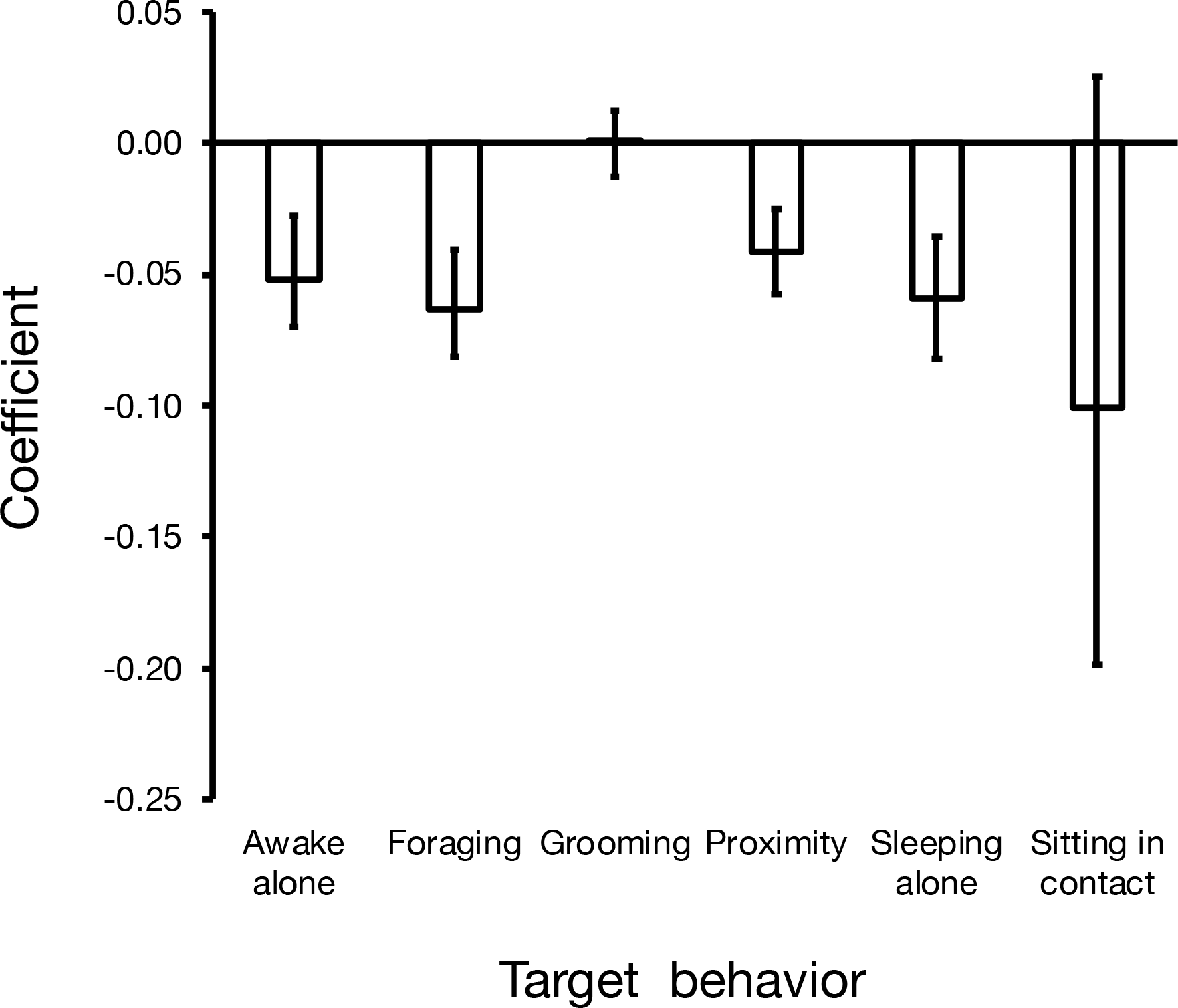
Effects of the target behavior on the relation between target rank and subject’s glance rate. Coefficients of the separate Poisson regressions (and 95% confidence intervals), controlling for target-subject kinship.

Focusing on grooming, we analyzed the effects of the characteristics of the grooming dyad on the glances the subject directed to it. Neither the rank difference between groomer and groomee (coeff.=0.010, z=0.75, N=1520, P=0.451), nor their degree of kinship (coeff.=0.206, z=0.97, N=1520, P=0.330), nor their baseline frequency of grooming (coeff.=−0.156, z=−0.11, N=1520, P=0.916) affected the glances they received from the subject.

## DISCUSSION

The results of this study show that the behavior of group mates is a potent modulator of the attention mandrills pay to them. Mandrills looked most often at foraging individuals and showed particular interest for individuals involved in social interactions. Mandrills also differentiated sleeping from awake individuals, showing that they were sensitive to rather subtle differences in the behavior of their group mates.

A previous study showed mandrills devoted particular attention at individuals that had recently been involved in a fight (individuals that are also known to be particularly likely to redirect aggression to bystanders; Schino & Marini, 2012; Schino & Sciarretta, 2016). In the same vein, in this study more glances were directed at high-ranking and at unrelated individuals (see also Emory, 1976; Pitcairns, 1976), and very little glances were directed at sleeping individuals (that presumably present very little danger). These observations are coherent with field studies reporting a greater attention devoted to potentially dangerous individuals and social contexts (e.g., Gaynor & Cords, 2012; Watts, 1998). Generally speaking, one of the functions of monitoring group mates seems to be the detection and possibly the anticipation of the risk of aggression, although in the field vigilance against predators often makes the identification of social monitoring more difficult (Allan & Hill, 2017; Treves, 2000).

It should be noted that our observation that mandrills looked at unrelated individuals more than at their kin stands in direct contrast with the results of a previous study on the same social group (Schino & Sciarretta, 2016). While it is difficult to find an explanation for this discrepancy, it should be noted that the two studies differed somewhat in the group composition (in the previous study several subadult, natal males were present), in the data collection procedure (in the previous study observation sessions had two targets instead of one, and the target activity was not controlled) and in the data analysis (in the previous study averages per dyad, rather than individual observation sessions, were the unit of analysis). None of these differences stands as an obvious candidate for explaining the different results obtained about the effect of kinship on glance rate. At any rate, the differences between the results of these two studies highlight that patterns of social attention may in some cases be sensitive to methodological details that appear of minor importance (see also Allan & Hill 2017). It should however also be noted that other patterns of social attention seem in contrast to be extremely robust even in the face of interspecific and methodological differences. In fact, our observation that mandrills looked at high-ranking individuals more than at low-ranking individuals is consistent with several previous studies conducted both by us and by others (Emory, 1976; Keverne et al., 1978; Pitcairns, 1976; Schino & Sciarretta, 2016).

Both primates and non-primates have been shown to be particularly attentive to foraging individuals (Ottoni et al., 2005; Range & Huber, 2007; Scheid et al., 2007). Our observation that mandrills paid particular attention to foraging individuals suggests that, even in a captive environment, group mates may constitute an important source of ecologically relevant information. Clearly, in captivity the degree to which foraging group mates attract attention is likely to depend on the details of the routine of food administration. When food is dispersed in the enclosure, group mates can more easily provide useful information than when food is clumped in a single pile or always put in the same place.

Mandrills seemed to be interested in the social interactions among their group mates. Very little comparable information is available in the literature, a notable exception being a study by Kret et al. (2016) that showed bonobos biased their attention towards pictures of grooming and sexual behavior. Whether this is a general primate pattern remains to be ascertained. Intriguingly, in mandrills grooming individuals were looked at irrespective of their dominance rank. While for most of the target's behaviors that we examined high-ranking individuals were looked at more than low-ranking individuals, grooming low-ranking individuals were looked at as much as grooming high-ranking individuals. Also, grooming was looked at irrespective of the rank difference between groomer and groomee, of their kinship, and (contrary to our expectations) of their baseline frequency of grooming (that is, rare grooming events were not looked at more than common events). Grooming seemed to be always equally interesting.

The general attention paid to grooming cannot be explained in terms of the need to monitor potentially dangerous situations, as grooming individuals do not constitute an impending threat. In contrast, the interest shown towards grooming individuals is likely to derive from the need to monitor and update the information about social relationships among group mates. Primates may need an unbiased estimate of the grooming interactions occurring among group mates and, accordingly, mandrills did not bias their attention towards any particular class of grooming dyads. It is also interesting to note that mandrills differentiated between grooming and sitting in contact. Although human observers often conflate grooming and passive contact as indicators of general affiliation (e.g., Silk et al., 2013), these two behaviors imply different costs and benefits and seem to be regarded by monkeys themselves as differentially worth of attention.

Visual monitoring has been shown to have a specific genetic basis (Watson et al., 2015) and is likely to have adaptive consequences by allowing the acquisition of valuable social and ecological knowledge. Primates are known to use in agonistic contexts the social knowledge they derive from monitoring group mates (Perry et al., 2004; Schino et al., 2006), although we still have very little evidence that this social knowledge actually has positive functional consequences (Tiddi et al., 2017). Elucidating the fitness consequences of variation in social knowledge seems thus to be an important research priority for future studies.

## ACKNOWLEDGEMENTS

We thank the Rome zoo (Bioparco) for allowing us to study their mandrill colony, and Giorgio Manzi for his support. Sarah Brosnan and two anonymous reviewers provided constructive criticism.

## DATA AVAILABILITY STATEMENT

Data available on request from the authors.

